# Anillin proteins stabilize the cytoplasmic bridge between the two primordial germ cells during *C. elegans* embryogenesis

**DOI:** 10.1101/126284

**Authors:** Eugénie Goupil, Rana Amini, Jean-Claude Labbé

## Abstract

Stable cytoplasmic bridges arise from failed cytokinesis, the last step of cell division, and are a key feature of syncytial architectures in the germ line of most metazoans. Whereas the *C. elegans* germ line is syncytial, its formation remains poorly understood. We studied the role of ANI-2, a noncanonical and shorter form of the actomyosin scaffold protein anillin that is expressed specifically during embryogenesis in the germ line precursor blastomere, P_4_. We found that the P_4_ blastomere does not complete abscission following cytokinesis, leaving a stable cytoplasmic bridge between the two daughter cells. Interestingly, depletion of ANI-2 results in a regression of the membrane partition between the two cells, indicating that ANI-2 is required to stabilize the cytoplasmic bridge. We identified several contractility regulators that, like ANI-2, localize to the cytoplasmic bridge and are required to stabilize it. Epistatic analysis of these regulators’ mutual dependencies revealed a pathway in which Rho regulators promote ANI-2 accumulation at the stable cytoplasmic bridge, which in turns promotes the accumulation of the non-muscle myosin II NMY-2 and the midbody component CYK-7 at the bridge, in part by limiting the accumulation of canonical anillin ANI-1. Our results uncover key steps in *C. elegans* germ line formation and define a set of conserved regulators that ensure the proper stability of the primordial germ cell cytoplasmic bridge during embryonic development.

## INTRODUCTION

Cytokinesis, the last step of cell division in which two daughter cells are physically separated, is a highly-coordinated event (reviewed in D’Avino et al., 2015; Green et al., 2012). It initiates during anaphase, when antiparallel microtubules in the mitotic spindle midzone organize signaling to the cell cortex to specify the future site of ingression, in part by recruitment of the centralspindlin complex (Mishima et al., 2002; Wheatley and Wang, 1996). This leads to the cortical recruitment and activation of the small GTPase Rho to the site of ingression, where it orchestrates the assembly and furrowing of a contractile ring composed of several proteins including actin filaments, non-muscle myosin II, septins and anillin (D’Avino et al., 2015; Green et al., 2012). Ingression of the contractile ring compacts the central portion of the mitotic spindle and generates a transient intercellular bridge termed midbody (Mullins and Biesele, 1977). As furrow constriction nears completion, the contractile ring becomes progressively tighter and matures into the midbody ring, a dense structure that forms around the center of the midbody. Midbody ring formation blocks the cytoplasmic exchange between the two daughter cells, effectively resulting in cytoplasmic isolation (Agromayor and Martin-Serrano, 2013; Green et al., 2013). Cytokinesis completes with cellular abscission, when recruitment of the endosomal sorting complexes required for transport (ESCRT) proteins promotes membrane fusion and midbody ring release, generating two physically separated daughter cells (Carlton et al., 2012; Green et al., 2013; Morita et al., 2007).

All cytokinetic steps are closely regulated to avoid cytokinetic failure, cellular binucleation and subsequent aneuploidy. In certain contexts, however, such as the germ line of many organisms across phyla, controlled cytokinesis failures take place during development, progressively leading to the formation of a cluster of joined cells that share cytoplasm through stable intercellular bridges, thus forming a syncytium (Fawcett et al., 1959; Greenbaum et al., 2007; Haglund et al., 2011). Such mechanism was studied in detail in mouse spermatocytes, in which the spermatocyte-specific protein TEX14 binds to the midbody ring protein CEP55 and inhibits recruitment of the ESCRT I protein TSG-101, thus effectively blocking the completion of abscission (Greenbaum et al., 2007; Greenbaum et al., 2006).Whether blocking abscission is a conserved feature of germ line formation is unclear.

The *C. elegans* adult germ line is organized as a syncytium in which each germ cell possesses a stable intercellular bridge that connects it to a central core of common cytoplasm, known as the rachis (Hubbard and Greenstein, 2005). The ring that stabilizes each germ cell intercellular bridge is enriched in regulators of actomyosin contractility that are also found in cytokinetic rings, including two forms of the actomyosin protein anillin (Amini et al., 2014; Maddox et al., 2005; Zhou et al., 2013). In other systems, anillin was shown to function as an adaptor protein that scaffolds contractility regulators to promote cytokinetic ring positioning and constriction (Hickson and O’Farrell, 2008; Oegema et al., 2000; Piekny and Glotzer, 2008). Anillin is proposed to achieve this in part by binding to the plasma membrane via its C-terminal Pleckstrin homology (PH) and C2 domains (Liu et al., 2012; Oegema et al., 2000; Sun et al., 2015), and interacting with actomyosin contractility regulators, such as RhoA, MgcRacGAP/RacGAP50 and Ect2, through its actin, myosin and anillin homology (AH) domains (D’Avino et al., 2008; Frenette et al., 2012; Gregory et al., 2008; Piekny and Glotzer, 2008; Straight et al., 2005). Disruption of anillin interaction with its partners leads to cytokinesis failure (Kechad et al., 2012; Oegema et al., 2000; Straight et al., 2005). In addition to canonical ANI-1, *C. elegans* stable germ cell intercellular bridges are enriched in a shorter, non-canonical anillin isoform, termed ANI-2, which lacks N-terminal putative actin and myosin binding domains, but retains the C-terminal AH and PH domains (Amini et al., 2014; Maddox et al., 2005). Animals depleted of ANI-2 display a disorganized germ line made of incomplete partitions comprising multiple nuclei (Amini et al., 2014; Green et al., 2011), indicating a role for ANI-2 in organization of stable germ cell intercellular bridges and the adult gonad architecture.

How the syncytial organization of the germ line arises during *C. elegans* development is not known. All germ cells originate from a single germ line precursor blastomere, termed P_4_, that arises after successive asymmetric divisions during embryogenesis (Wang and Seydoux, 2013). Around the 100-cell stage, P_4_ divides into the two primordial germ cells (PGCs), Z_2_ and Z_3_, which remain mitotically quiescent throughout embryogenesis and initiate proliferation after hatching, at the mid-L1 larval stage, to give rise to the entire syncytial germ line (Hirsh et al., 1976; Sulston et al., 1983). Interestingly, unlike ANI-1 that is present in all cells, ANI-2 is specifically found in the P_4_ blastomere during embryogenesis, localizes to the nascent furrow during P_4_ division and is enriched at the midbody ring between Z_2_ and Z_3_ after cytokinesis (Amini et al., 2014). Whether ANI-2 accumulation in the two primordial germ cells during embryogenesis is functionally relevant for proper adult germ line formation was unknown. Here we demonstrate that the P_4_ blastomere initiates cytokinesis but does not complete abscission, resulting in a stable cytoplasmic bridge between the two PGCs. We show that ANI-2 accumulation at the cytoplasmic bridge, along with several other contractility regulators, is essential to ensure its stability throughout embryogenesis and give rise to a fully organized and functional adult germ line.

## RESULTS

### ANI-2 regulates *C. elegans* primordial germ cell formation

*ani-2*(-) mutant animals develop into sterile adults, with their germ cells becoming increasingly multinucleated starting from late larval stages, however the ANI-2 protein is already present during embryogenesis, upon birth of the germ line precursor blastomere, P_4_ (Amini et al., 2014). To find out whether this embryonic pool of ANI-2 plays an earlier role in germ line development, we used RNAi to deplete ANI-2 from animals expressing fluorescent markers enabling the visualization of plasma membrane (mNeonGreen fused to the PH domain of PLCδ, hereafter mNG::PH) and cell nuclei (mCherry fused to histone H2B, hereafter mCherry::H2B; Fig. 1A). Live imaging of ANI-2-depleted embryos revealed that they proceeded through early development normally and that, as in controls, P_4_ divided at the ∼100-cell stage (Fig. 1A and B). Interestingly however, the membrane partition separating Z_2_ and Z_3_ eventually regressed in 49% of ANI-2-depleted embryos (n=43), on average 137 ± 23 min after the membrane had visibly completed its ingression (Fig. 1A and C, Movie S1). Regression occurred on average at the 215 ± 36-cell stage (Fig. 1B), near the end of gastrulation, when Z_2_ and Z_3_ typically undergo rotation in the long axis of the embryo (Chisholm and Hardin, 2005). This phenotype is reminiscent of the previously reported membrane collapse observed in the germ cells of adult animals lacking ANI-2 (Amini et al., 2014). RNAi-mediated depletion of ANI-2 was effective as judged by the strong decrease in ANI-2 signal in immunofluorescence experiments (Fig. 1D). Accordingly, *ani-*2(RNAi) embryos grew into sterile adults with a disorganized germ line architecture (Fig. 1E). These results indicate that ANI-2 is not essential for cytokinetic furrow ingression of the P_4_ germ line precursor blastomere but is required to ensure proper completion of cytokinesis and formation of the two PGCs.

**Figure 1.**
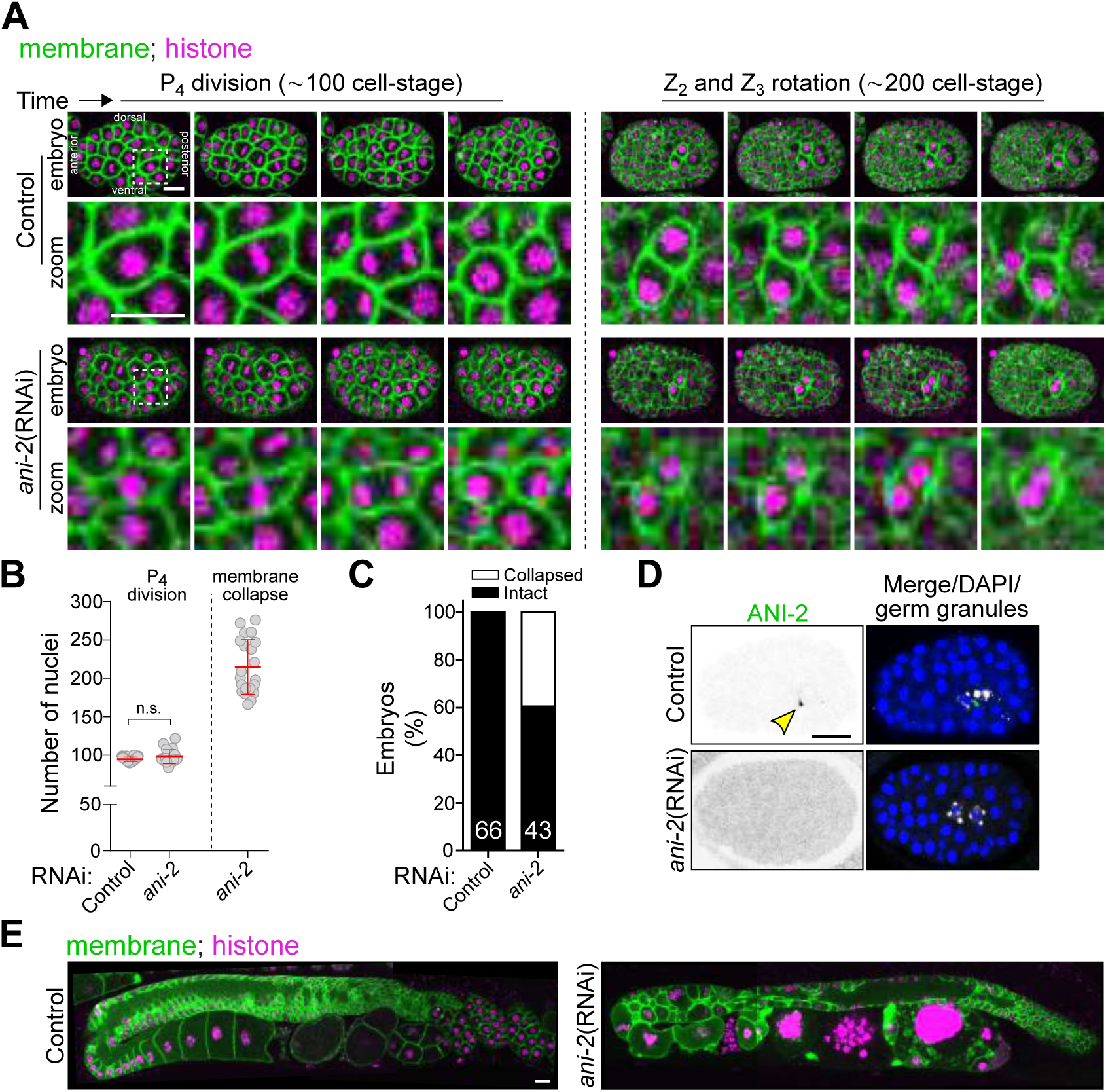
Depletion of ANI-2 causes membrane collapse between the two primordial germ cells. (A) Confocal images (maximum intensity projections of 2 consecutive Z stacks) of control and *ani-*2(RNAi) embryos expressing membrane (mNG::PH, green) and histone (mCherry::H2B, magenta) markers during birth of the two PGCs. The region delineated by the white dashed square around P_4_ is magnified by 3.5-fold in the inset. (B) Quantification of the number of nuclei upon P_4_ division and membrane collapse in control and *ani-2*(RNAi) embryos. The red bars represent average ± SD. n.s. = p>0.05. (C) Percentage of collapsed partitions between the two PGCs in control and *ani-2*(RNAi) embryos. (D) Mid-section confocal images of control and *ani-2*(RNAi) embryos fixed and stained with antibodies against ANI-2 (green), germ granules (white) and DAPI (blue). Yellow arrowhead points to the ANI-2 focus in the control. (E) Mid-section confocal images of the adult germ line in the progeny (F1) of control and *ani-2*(RNAi) animals. Scale bars, 10 μm.

### ANI-2 stabilizes the cytoplasmic bridge between the two PGCs during embryogenesis

ANI-2 could either function in the P_4_ cell to ensure completion of cytokinesis, or it could act to stabilize the cytoplasmic bridge after a developmentally-programmed incomplete cytokinesis. To discriminate between these possibilities, we set out to determine whether the P_4_ blastomere undergoes complete cytokinesis. This was done by monitoring the localization of GFP-tagged non-muscle myosin II (NMY-2::GFP) in embryos co-expressing fluorescently tagged markers for the plasma membrane (TagRFP::PH) and germ cells (PGL-1::RFP), and comparing events occurring in the dividing P_4_ blastomere with those of neighboring somatic cells (Fig. 2A-C and S1). Cytokinesis was previously shown to occur in four successive phases: contractile ring assembly, contractile ring ingression, cytoplasmic isolation and cellular abscission (D’Avino et al., 2015; Green et al., 2012). We found that the first three phases of cytokinesis were normal in the P_4_ blastomere and that it underwent proper cytoplasmic isolation in all embryos (n=27; Fig. 2C and Movie S2), although ingression dynamics were slightly but significantly delayed compared to somatic cells (Fig. 2D-E). Depletion of ANI-2 did not perturb ingression dynamics in either P_4_ or somatic cells (Fig. 2F). This indicates that the first three phases of cytokinesis occur normally in the P_4_ blastomere and that they do not critically rely on ANI-2 activity.

**Figure 2.**
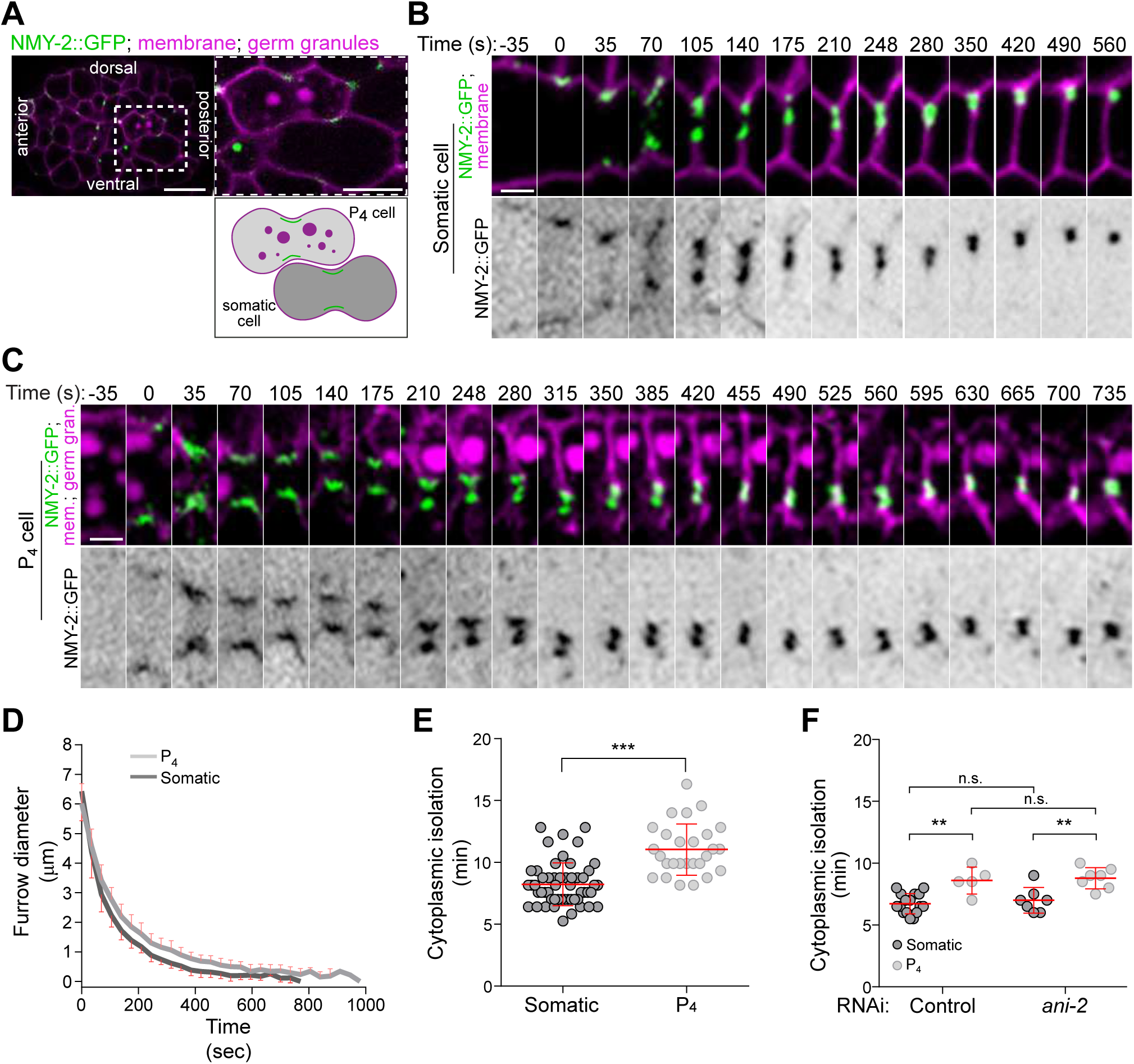
The dividing P_4_ blastomere undergoes proper cytoplasmic isolation. (A) Confocal images (maximum intensity projection of 3 consecutive Z planes) of a control embryo expressing NMY-2::GFP (green), a membrane marker (TagRFP::PH, magenta) and a germ granule marker (PGL-1::RFP, magenta). The region delineated by the white dashed square is magnified by 4-fold in inset and shows a somatic cell and a P_4_ cell upon initiation of cytokinetic furrow ingression. The schematic illustration depicts the region of interest for monitoring furrow progression. (B, C) Time-lapse confocal images of the division plane in a somatic (B) and P_4_ (C) cell from a control embryo expressing the markers described in (A). Time 0 (s) corresponds to the onset of cytokinetic furrow initiation. (D-E) Quantification of cytokinetic furrow diameter (D) and duration of cytokinetic furrow ingression (E) from the onset of furrow initiation to cytoplasmic isolation in wild-type somatic (n= 54, dark grey) and P_4_ (n=27, light grey) cells. (F) Quantification of the duration of cytokinetic furrow ingression from the onset of furrow initiation to cytoplasmic isolation in somatic (dark grey) and P_4_ (light grey) cells of control or *ani-2*(RNAi) embryos. The red bars represent average ± SD. **p<0.01, ***p<0.001, n.s. = p>0.05. Scale bar for full embryo, 10 μm; scale bar for insets, 2 μm.

We then monitored cellular abscission, the last phase of cytokinesis, by measuring the timing of midbody ring-associated NMY-2::GFP release from the interstitial membrane separating two sister cells (Fig. S1D-F). In somatic cells, we found that the midbody ring took on average 14.3 ± 5.1 min to dissociate from the interstitial membrane separating the two sister cells (n=35; Fig. 3A-B). In sharp contrast, the NMY-2::GFP signal remained present at the Z_2_/Z_3_ interstitial boundary, even up to 200 min after cytoplasmic isolation (n=30; Fig 3A-B and Movie S2), and its intensity was stable over this entire period (Fig. 3D). Similar results were obtained upon monitoring the localization of GFP-tagged versions of the canonical anillin ANI-1, the septin UNC-59 and the midbody-associated protein CYK-7 (Fig. 3C and D). These results indicate that the *C. elegans* P_4_ blastomere fails to complete cellular abscission, resulting in the two PGCs maintaining a stable cytoplasmic bridge.

**Figure 3.**
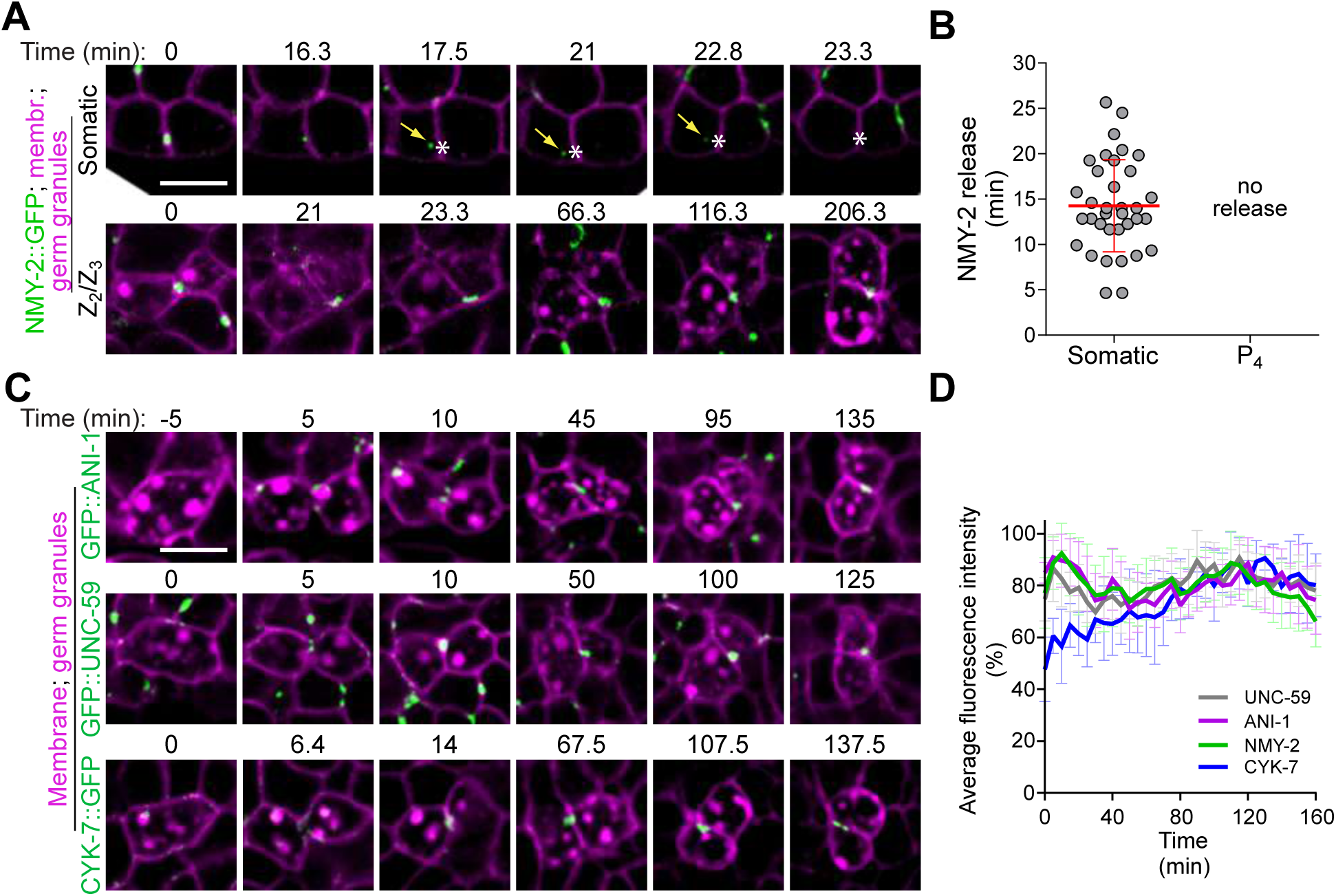
The P_4_ blastomere does not complete cellular abscission. (A) Mid-section confocal time-lapse images of sister somatic cells (top panels) and PGCs (bottom panels) from a control embryo expressing NMY-2::GFP (green), a membrane marker (TagRFP::PH, magenta) and a germ granule marker (PGL-1::RFP, magenta). Time 0 corresponds to cytoplasmic isolation. Yellow arrows point to the midbody ring-associated NMY-2 focus upon release in the cytoplasm and white asterisks indicate the position in the membrane from which the midbody ring was released (top panels only). (B) Quantification of the time of midbody ring release relative to the time of cytoplasmic isolation in control somatic and germ cell siblings. The red bar represents average ± SD. (C) Mid-section confocal time-lapse images of control germ cells expressing membrane and germ granule markers (both in magenta) and one of three midbody ring markers (GFP::ANI-1, GFP::UNC-59 or CYK-7::GFP, green). Time 0 corresponds to cytoplasmic isolation. (D) Quantification of the average fluorescence intensity (± SD) of the unreleased NMY-2::GFP, GFP::ANI-1, GFP::UNC-59 and CYK-7::GFP foci at the stable cytoplasmic bridge relative to the time of cytoplasmic isolation in control PGCs. Scale bar, 5 μm.

We previously found that ANI-2 accumulates specifically in the P_4_ blastomere and between the two PGCs (Amini et al., 2014), however it remained unclear whether ANI-2 was continuously present at the stable cytoplasmic bridge. We used indirect immunofluorescence to carefully monitor the distribution of ANI-2 in the embryo, from the division of P_4_ at the ∼100-cell stage to the onset of epithelial morphogenesis 150 minutes later, at the ∼350-cell stage. As previously reported (Amini et al., 2014), we found that ANI-2 was specifically present at the cortex of P_4_ and became enriched at the cleavage furrow upon cytokinesis (Fig. 4A). ANI-2 accumulated at the midbody ring between Z_2_ and Z_3_ and remained visible as a focus between the two PGCs up to the ∼350-cell stage (Fig. 4A). Quantitative analysis of the ANI-2 signal fluorescence intensity and distribution revealed that the volume of the ANI-2 focus increased slightly but steadily from its formation, upon P_4_ division, until it had nearly doubled in size at the ∼350-cell stage (Fig. 4B-C). To assess whether the ANI-2 focus undergoes shape change during this time we used the well-defined sphericity index, which reports on an object’s surface area relative to its volume (see Methods for details; Wadell, 1935). We found that the sphericity value of the ANI-2 focus remained constant (Fig. 4C), indicating that the increase in ANI-2 focus volume is not accompanied by a significant change in focus shape. ANI-2 colocalized and behaved similarly to GFP-tagged versions of the non-muscle myosin II NMY-2 and the midbody protein CYK-7 (Fig. S2), the dynamics of which can be monitored in real time during embryogenesis. These results indicate that ANI-2 is a stable component of the PGC cytoplasmic bridge, from its formation upon P_4_ cytokinesis to at least the onset of epithelial morphogenesis at the ∼350-cell stage. Importantly, these results indicate that ANI-2 is not required to ensure P_4_ cytokinetic completion but is necessary to stabilize the cytoplasmic bridge between Z_2_ and Z_3_ at the ∼200-cell stage of embryogenesis.

**Figure 4.**
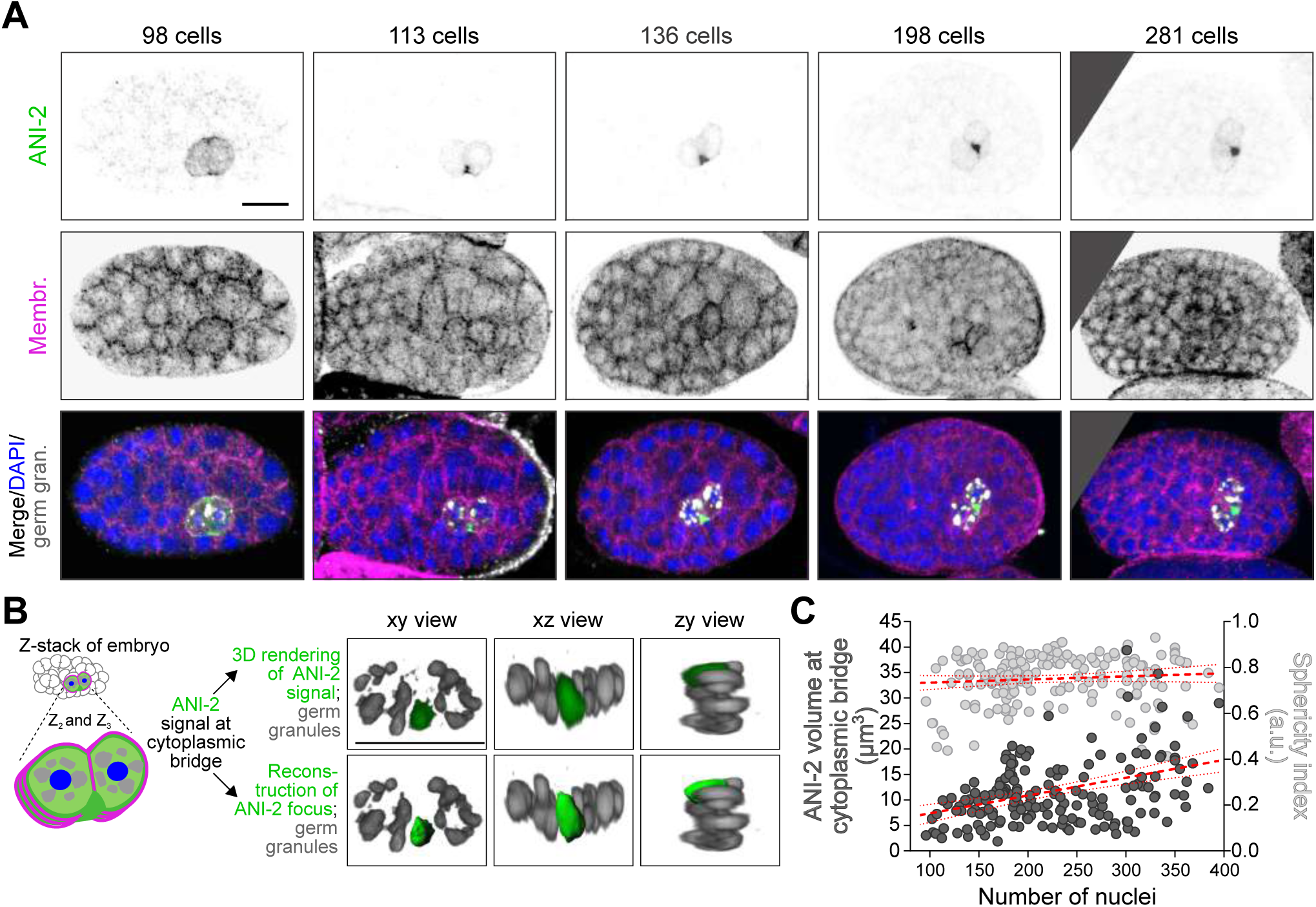
ANI-2 is a stable component of the PGC cytoplasmic bridge. (A) Confocal images (maximum intensity projections of 5-7 consecutive Z planes) of control embryos between the ∼100- and ∼300-cell stages expressing a membrane marker (GFP::PH), and fixed and stained with antibodies against GFP (magenta), ANI-2 (green), germ granules (white) and DAPI (blue). (B) Schematic representation of ANI-2 localization and depiction of the method used to measure the ANI-2 focus volume at the cytoplasmic bridge between the two PGCs. The 3D rendering of the original ANI-2 fluorescence signal obtained by immunofluorescence (green) and the reconstruction of the focus using Imaris (green) with the germ granules (grey) are shown (see Methods for details). (C) Quantification of the ANI-2 focus volume (dark grey, left axis) and sphericity (light grey, right axis) at the PGC cytoplasmic bridge as a function of number of nuclei in control embryos. Linear regressions (full red line) and confidence intervals (dashed red line) are shown above each data set. Scale bars, 10 μm.

### Several contractility regulators stabilize the cytoplasmic bridge between the two PGCs

To better understand how ANI-2 stabilizes the PGC cytoplasmic bridge, we next sought to identify other regulators of membrane stabilization amongst genes encoding cytokinetic regulators and/or genes whose depletion was previously reported to phenocopy the loss of ANI-2 function in the adult germ line (22 genes, see Table S2 for a complete list; Green et al., 2011). We used partial RNAi-mediated depletion conditions for each gene, as the complete depletion of many of these would cause a failure in the first embryonic division, thus precluding a study of Z_2_ and Z_3_ later in embryogenesis. For each gene tested, our depletion regime resulted in a small number of binucleated cells and a high percentage of embryonic lethality, but yet allowed the division of P_4_ and birth of the two PGCs (Fig. S3A; see Methods for details).

Live imaging of embryos depleted of the different genes of interest revealed that in all cases in which the initial steps of P_4_ blastomere cytokinesis were not compromised, P_4_ underwent cytoplasmic isolation at the ∼100-cell stage, as in control. However, we observed a variable proportion of embryos in which the membrane between Z_2_ and Z_3_ had regressed after depletion of 12 of the 22 genes, a phenotype similar to that observed after ANI-2 depletion (Fig. 5A). These genes encode the small GTPase RHO-1 and its guanine-nucleotide exchange factor ECT-2, the two centralspindlin subunits CYK-4 and ZEN-4 (Mishima et al., 2002), the non-muscle myosin II NMY-2, the myosin chaperone UNC-45 (Kachur et al., 2008), the actin subunit ACT-4, the formin CYK-1, the canonical anillin ANI-1, the midbody-associated component CYK-7 (Green et al., 2011), the non-canonical Patched receptor PTC-2 (Kuwabara et al., 2000), and F30B5.4, a putative homolog of the mammalian oxidative stress-induced growth inhibitors, OSGIN1/2 (Huynh et al., 2001). The proportion of embryos that displayed a membrane regression phenotype varied depending on the gene depleted (Fig. 5B). Likewise, while most membranes collapsed near the embryonic stage where collapse was observed in ANI-2-depleted embryos, some collapse events occurred at later stages (Fig. 5C). These variations could reflect different requirements for these proteins in regulating membrane stability but are more likely a consequence of partial RNAi-mediated depletions in each case. Insufficient depletion could also account for the absence of membrane collapse in 10 of the 22 regulators tested, although expected phenotypes were observed in each case, including defects in embryonic viability (Fig. S3A) and cytokinesis (Fig. S3B). Taken together, these results highlight a set of regulators that are required to stabilize the cytoplasmic bridge between the two PGCs during *C. elegans* embryogenesis.

**Figure 5.**
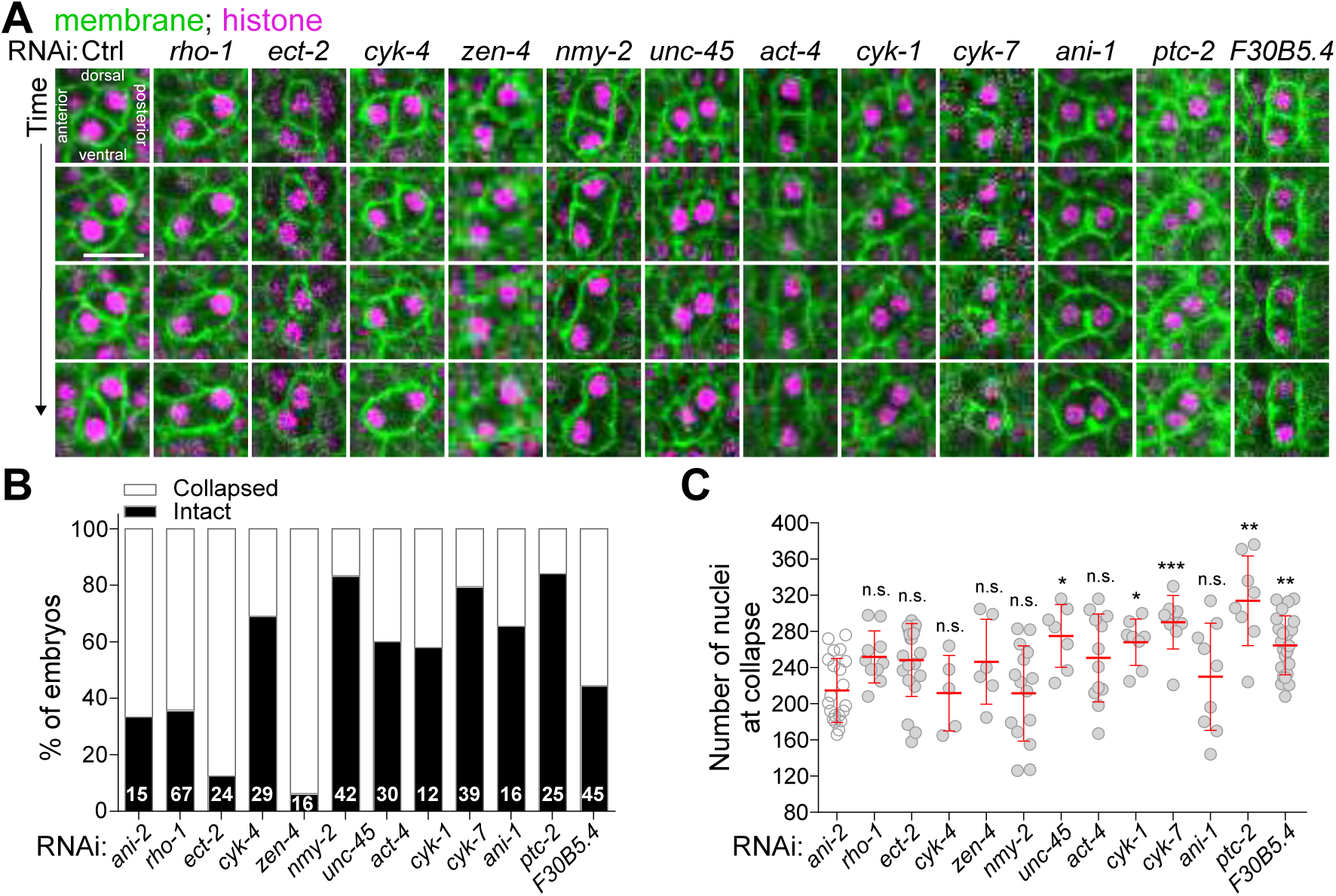
Several contractility regulators are required to stabilize the PGC cytoplasmic bridge. (A) Mid-section confocal time-lapse images of PGCs from embryos expressing membrane (mNG::PH, green) and histone (mCherry::H2B, magenta) markers. Animals were differentially depleted of contractility regulators by RNAi (see Table S2) and stability of the PGC cytoplasmic bridge was assessed by monitoring membrane signal. Scale bar, 10 μm. (B-C) Quantification of the percentage of embryos undergoing membrane collapse (B) and number of nuclei at membrane collapse (C) for each condition. The red bars represent average ± SD.

### The Rho pathway promotes ANI-2 localization at the stable PGC cytoplasmic bridge

We then sought to determine whether regulators of membrane stability between the two PGCs function with ANI-2 in this process. Hermaphrodite animals were partially or strongly depleted of the different proteins of interest by RNAi (as above) and the volume of the ANI-2 focus between Z_2_ and Z_3_ was measured in embryos containing 150-250 nuclei, the stage at which the membrane between PGCs receded in *ani-2*(RNAi) embryos. As expected, depleting ANI-2 by RNAi resulted in a strong reduction of the ANI-2 signal and focus volume compared to control (Fig. 6A and B), providing a comparative baseline for this assay. Among the proteins required to maintain a stable PGC cytoplasmic bridge, we found that depletion of five of them resulted in a significant decrease in the volume of the ANI-2 focus: the two centralspindlin subunits CYK-4 and ZEN-4, RHO-1, ECT-2 and F30B5.4 (Fig. 6A and B). Mispositioning of the ANI-2 focus at cortical regions other than the Z_2_/Z_3_ interstitial boundary was observed in ∼30% of embryos depleted of RHO-1 or CYK-4, and ∼7% of ECT-2-depleted embryos. These results suggested that ANI-2 could function downstream of Rho signaling in this process. To test this hypothesis, we analyzed the localization of CYK-4::GFP in embryos depleted of ANI-2 and other contractility regulators. We found that CYK-4::GFP co-localizes with ANI-2 at the stable PGC cytoplasmic bridge in control embryos, and that CYK-4::GFP accumulation at the bridge was not perturbed by depletion of ANI-2 or any of the other contractility regulators herein tested (Fig. S4A and B). These results indicate that Rho pathway regulators and F30B5.4 function upstream of ANI-2 to promote its localization at the stable PGC cytoplasmic bridge.

**Figure 6.**
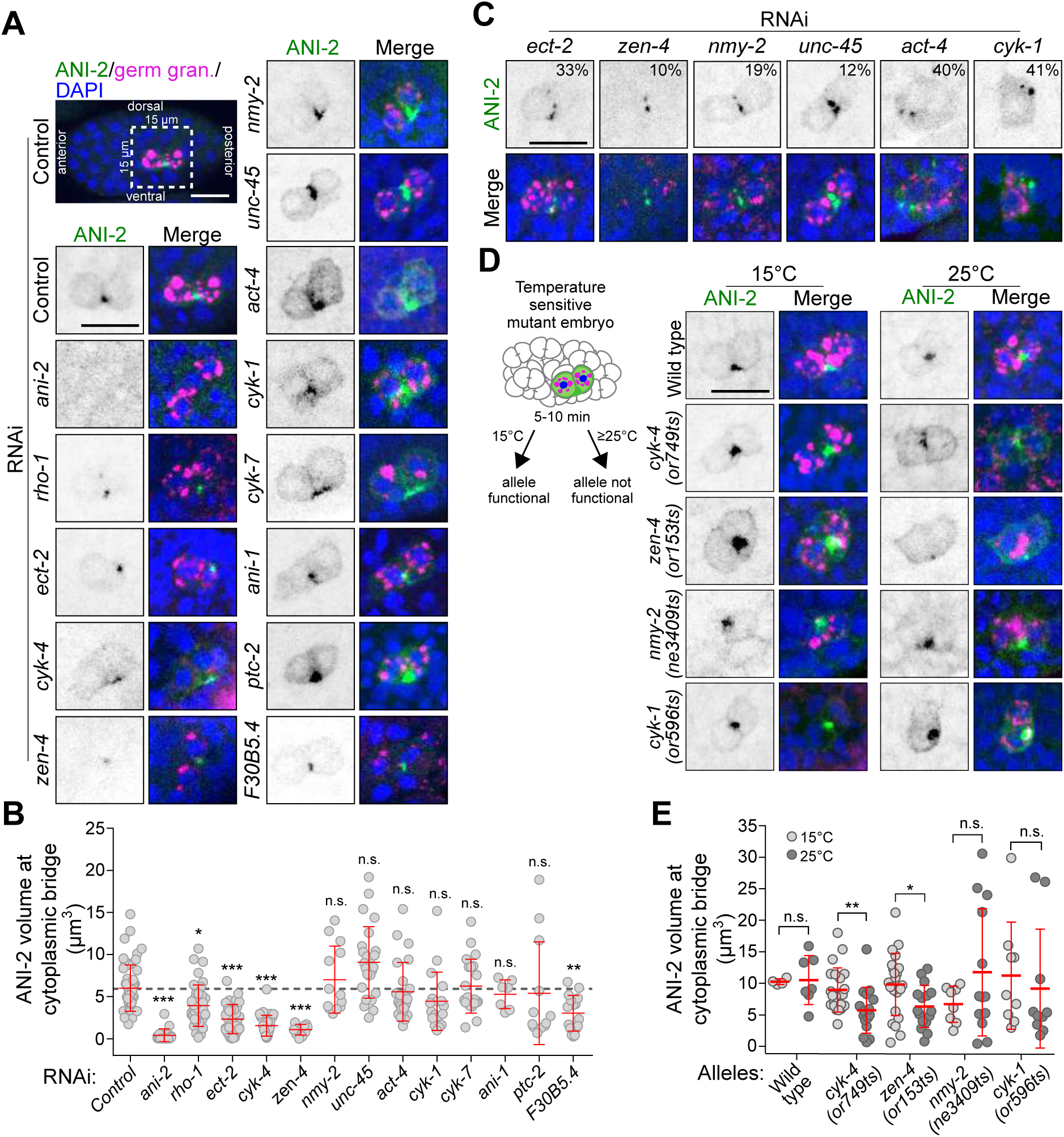
Rho pathway regulators promote ANI-2 accumulation at the stable PGC cytoplasmic bridge. (A, C, D) Confocal images (maximum intensity projections of 5-7 consecutive Z planes) of PGCs in 150-250-cell stage embryos, fixed and stained with antibodies against ANI-2 (green), germ granules (magenta) and DAPI (blue). The insets depict PGCs from embryos after RNAi depletion of the genes indicated (A, C) or PGCs from embryos bearing fast-acting, temperature-sensitive alleles of the genes indicated and acquired after continuous growth at permissive temperature (15°C) or after a 5-10-minute upshift at restrictive temperature (25°C), as depicted in the schematic (D). In (C), the number (%) represents the proportion of embryos displaying fragmentation of the ANI-2 focus. Scale bars, 10 μm. (B, E) Quantification of the focus volume of ANI-2 at the PGC cytoplasmic bridge in embryos depleted of the indicated genes by RNAi (B) or in embryos bearing the temperature-sensitive alleles of the indicated genes and maintained at permissive (15°C) or restrictive (25°C) temperature (E). The red bars represent average ± SD. *p<0.05, **p<0.01, ***p<0.001, n.s. = p>0.05.

In addition, depletion of ECT-2, ZEN-4, NMY-2, UNC-45, ACT-4 and CYK-1 lead to a variable proportion of embryos displaying fragmentation of the ANI-2 signal into multiple smaller foci with a combined volume indistinguishable from that of control embryos (Fig. 6C and S3E). These defects in ANI-2 organization are likely a consequence of Z_2_/Z_3_ interstitial membrane collapse, since they are more frequent in conditions that induce the highest frequency of collapse. Depletion of ANI-1, CYK-7, PTC-2 or any of the 10 proteins that did not cause membrane collapse had no effect on the accumulation of ANI-2 at the cytoplasmic bridge or its organization into a proper focus (Fig. 6A-B and S3C-D). These results indicate that most proteins whose depletion phenocopies that of ANI-2 in PGCs regulate the accumulation and/or organization of ANI-2 at the stable PGC cytoplasmic bridge.

To validate the results obtained by RNAi depletion, we took advantage of available mutants bearing fast-acting, temperature-sensitive (ts) alleles for *cyk-4, zen-4, nmy-2* and *cyk-1* (Davies et al., 2014). These mutant alleles provided us with the means to rapidly inactivate gene products at precise times, after the birth of the two PGCs, and thus exclude the possible confounding effects of early phenotypic alterations that remain a constant drawback of RNAi-mediated depletions. Embryos were upshifted to the restrictive temperature for 5-10 minutes before fixation and staining (Fig. 6D; see Methods), and analysis of the ANI-2 focus volume was performed on fixed 150-250-cell stage embryos, as above. In all mutant backgrounds, most embryos contained a few binucleated somatic cells, thus validating that the temperature upshift caused gene product inactivation. While temperature upshift did not change the volume of the ANI-2 focus in wild-type embryos, the same treatment caused a significant reduction of the ANI-2 focus volume in both *cyk-4*(*ts*) and *zen-4*(*ts*) mutant embryos, similar to what had been observed after RNAi depletion of these two genes (Fig. 6D and E). Upshifting *nmy-2*(*ts*) and *cyk-1*(*ts*) embryos did not change the ANI-2 focus volume, consistent with RNAi results (Fig. 6D and E). Together, these results are in agreement with those obtained by RNAi and indicate that genes encoding several contractility regulators, and particularly members of the Rho pathway, function with ANI-2 to regulate membrane stability between the two *C. elegans* PGCs.

### ANI-2 regulates the localization of NMY-2, CYK-7 and ANI-1 at the stable PGC cytoplasmic bridge

While Rho pathway regulators promote the maintenance of ANI-2 at the stable PGC cytoplasmic bridge, several contractility regulators affect the stability of this bridge without impacting ANI-2 localization, raising the possibility that they could function downstream of ANI-2. To assess this, we first looked at the localization of NMY-2 and CYK-7, two contractility regulators that colocalize with ANI-2 at the stable cytoplasmic bridge (Fig. 7A and C). We found that depletion of ANI-2 by RNAi significantly reduced the focus volume of both NMY-2::GFP and CYK-7::GFP at the stable cytoplasmic bridge between Z_2_ and Z_3_ (Fig. 7A-D). Similar results were obtained after RNAi depletion of the proteins required to localize ANI-2 at the stable bridge (Fig. 7A-D), consistent with these acting upstream of ANI-2 in the pathway. Interestingly, while depletion of NMY-2 caused a reduction of CYK-7 volume at the stable bridge (Fig. 7C and D), depleting CYK-7 did not affect NMY-2 localization (Fig. 7A and B), suggesting that NMY-2 acts upstream of CYK-7 in this pathway. These results suggest that ANI-2 promotes the localization of NMY-2 at stable PGC cytoplasmic bridges and that, in turn, this enables the localization of CYK-7 at this structure.

**Figure 7.**
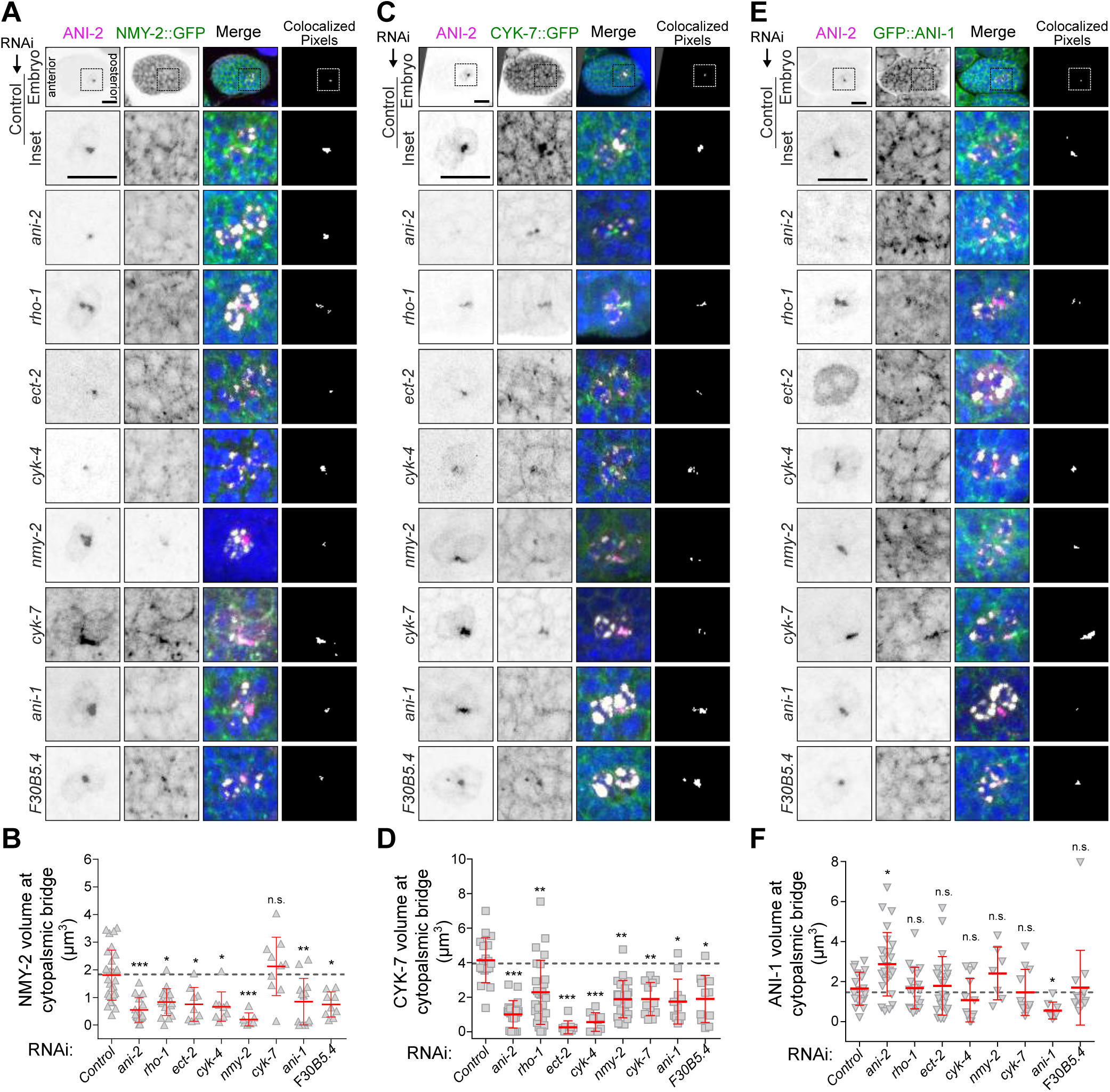
ANI-2 regulates the localization of NMY-2, CYK-7 and ANI-1 at the stable PGC cytoplasmic bridge. (A, C, E) Confocal images (maximum intensity projections of 5-7 consecutive Z planes) of PGCs in embryos at the 150-250-cell stage expressing either NMY-2::GFP (A), CYK-7::GFP (C) or GFP::ANI-1 (E), and fixed and stained with antibodies against GFP (green), ANI-2 (magenta), germ granules (white) and DAPI (blue). Embryos were depleted of the genes indicated by RNAi. Scale bar, 10 μm. (B, D, F) Quantification of the focus volume of NMY-2::GFP (B), CYK-7::GFP (D) or GFP::ANI-1 (F) at the PGC cytoplasmic bridge in embryos depleted of the indicated genes by RNAi. The red bars represent average ± SD. *p<0.05, **p<0.01, ***p<0.001, n.s. = p>0.05.

We and others previously demonstrated that the two *C. elegans* anillin proteins, the canonical ANI-1 and the short ANI-2, function opposite to each other in regulation of actomyosin contractility, in both the germ line and the 1-cell stage embryo (Amini et al., 2014; Chartier et al., 2011; Pacquelet et al., 2015; Rehain-Bell et al., 2017). We therefore asked whether this antagonistic relationship was likewise required for PGC cytoplasmic bridge stability. We found that while ANI-1 is required to maintain a stable partition between the two PGCs (Fig. 5), it is not required for CYK-4 or ANI-2 localization at the stable PGC cytoplasmic bridge (Fig. 6A-B and S4A-B), suggesting that it acts downstream of ANI-2 in this pathway. Depletion of ANI-1 caused a decrease of both NMY-2::GFP and CYK-7::GFP focus volumes at the stable bridge, indicating that ANI-1 acts upstream of these two regulators in the pathway (Fig. 7A-D). Consistent with this, we found that GFP::ANI-1 accumulates at the stable PGC cytoplasmic bridge and that its accumulation is independent of NMY-2 and CYK-7 (Fig. 7E and F). Interestingly, depletion of ANI-2 resulted in an increase in the GFP::ANI-1 focus volume at the stable bridge, indicating that ANI-2 acts to limit the amount of ANI-1 at this structure (Fig. 7F). Strikingly however, depleting Rho pathway regulators (CYK-4, ZEN-4, RHO-1, ECT-2) or F30B5.4, which are required to promote ANI-2 accumulation at the stable bridge (Fig. 6A and B), had no effect on GFP::ANI-1 accumulation (Fig. 7E and F). These results suggest that Rho pathway activity promotes the localization of both ANI-1 and ANI-2 at the stable PGC cytoplasmic bridge and that ANI-2 acts locally to limit the amount of ANI-1 at this structure, while actively promoting the accumulation of NMY-2 and CYK-7.

## DISCUSSION

In this study, we show that the embryonic germ line precursor cell P_4_ does not complete abscission upon cytokinesis and that the two primordial germ cells, Z_2_ and Z_3_, remain connected via a stable cytoplasmic bridge. We find that the short anillin ANI-2 and several actomyosin contractility regulators stably localize to the PGC cytoplasmic bridge and that the presence of ANI-2 at the Z_2_/Z_3_ interstitial boundary is not required to block abscission but is necessary to stabilize the bridge. Our results support a model in which Rho pathway regulators promote the loading of ANI-2 at the stable PGC cytoplasmic bridge, which in turns promotes the maintenance of NMY-2 and CYK-7 at the cytoplasmic bridge to ensure its stability (Fig. 8).

**Figure 8.**
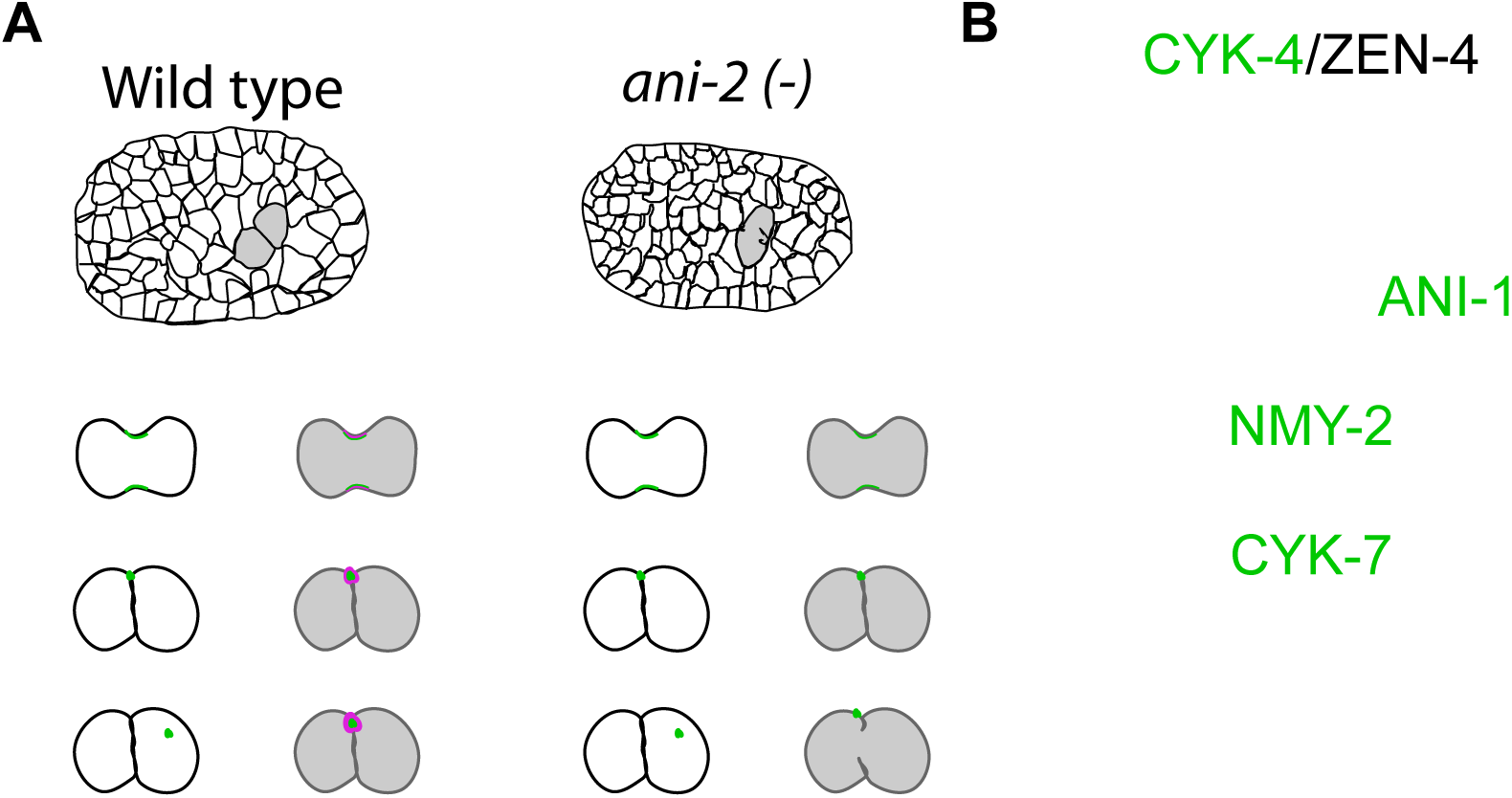
Regulation of PGC cytoplasmic bridge stability by actomyosin contractility regulators. (A) While somatic cells undergo proper midbody ring release (green) and cellular abscission in wild-type embryos, the P_4_ blastomere fails to complete abscission and stably maintains several contractility regulators, including ANI-2 (magenta), at the PGC cytoplasmic bridge. Depletion of ANI-2 has no effect on somatic cells but results in a destabilization of the PGC cytoplasmic bridge and regression of the membrane separating the two PGCs. (B) Working model depicting the proposed functional requirements for several contractility regulators in stabilizing the cytoplasmic bridge between the two PGCs during *C. elegans* embryogenesis. ANI-2 (magenta), ANI-1, CYK-4, NMY-2 and CYK-7 (green) all stably localize to the cytoplasmic bridge. While ANI-2 functions upstream of ANI-1 to limit its accumulation at the stable cytoplasmic bridge, it is unclear whether it regulates NMY-2 and CYK-7 localization directly (dashed line) or through ANI-1. See main text for details.

Our results suggest that the centralspindlin complex proteins CYK-4 and ZEN-4 are at the top of this hierarchy since depletion of either of them results in a membrane collapse and no other regulator perturbs the localization of CYK-4 at the stable cytoplasmic bridge. This contrasts with a previous study proposing that ANI-2 acts upstream of CYK-4 to regulate the stability of intercellular bridges in adult *C. elegans* germ cells (Zhou et al., 2013). This apparent discrepancy could originate from differences in ANI-2 requirements between embryos and adult animals or differences in RNAi depletion regimes, which were longer for depletions done in adult animals and may have led to cumulative defects and more severe germ line disorganization. The requirement of Rho regulators in ANI-2 localization at the cytoplasmic bridge is consistent with ANI-2 containing C-terminal domains that, in anillin proteins of other species, were reported to bind to RhoA/RHO-1, MgcRacGAP/CYK-4 and the plasma membrane (D’Avino et al., 2008; Gregory et al., 2008; Liu et al., 2012; Maddox et al., 2005). Depleting F30B5.4 results in similar defects as RHO-1 or ECT-2 depletion, however we currently do not know whether this regulator impinges on Rho signalling itself or where it acts between CYK-4 and ANI-2. Likewise, we do not know precisely where or how the other regulators of cytoplasmic bridge stability herein identified (UNC-45, PTC-2, CYK-1, ACT-4) act in this pathway, although work in *C. elegans* and other systems allows us to speculate that the myosin chaperone UNC-45 works with NMY-2 (Kachur et al., 2008) and that the formin CYK-1 and actin ACT-4 function together, downstream of Rho and partly in parallel of the other regulators (D’Avino et al., 2015; Green et al., 2012).

Previous studies carried out in the 1-cell stage embryo and adult gonad revealed that the canonical *C. elegans* anillin ANI-1 and the short anillin ANI-2 have opposing activities on actomyosin contractility (Amini et al., 2014; Chartier et al., 2011; Pacquelet et al., 2015; Rehain-Bell et al., 2017). Our work here reveals that both ANI-1 and ANI-2 are required to localize NMY-2 and CYK-7 at the PGC cytoplasmic bridge and maintain its stability. The N-terminal region of ANI-1 contains putative actin- and myosin-binding domains that could enable ANI-1 to bind NMY-2 directly and thus mediate NMY-2 localization. While these domains are not readily identified in ANI-2 (Maddox et al., 2005), we cannot exclude the possibility that ANI-2 could also directly interact with NMY-2 to affect its localization. However, whereas ANI-1 depletion has no noticeable effect on ANI-2 localization, depletion of ANI-2 results in an increase in ANI-1 accumulation at the stable cytoplasmic bridge, suggesting that ANI-2 acts upstream of ANI-1 in this cellular context. Another possibility is therefore that ANI-2 exerts its effect on NMY-2 entirely through ANI-1 recruitment at the stable cytoplasmic bridge, a scenario in which both a decrease or an increase in ANI-1 levels would impede NMY-2 localization. Interestingly, ANI-1 accumulation at the stable bridge did not vary upon depletion of the upstream Rho pathway regulators that are required for ANI-2 accumulation, suggesting that they promote the localization of ANI-1 at the stable cytoplasmic bridge independently of their activity on ANI-2 and that ANI-1 accumulation at the stable bridge is balanced by these two opposing activities. A Rho-dependent requirement for ANI-1 recruitment is consistent with the previously described function for Rho regulators acting upstream of anillin and enabling its cortical recruitment in other systems (Hickson and O’Farrell, 2008; Piekny and Glotzer, 2008). Alternatively, more complex molecular interactions between Rho regulators and *C. elegans* anillin proteins may impinge on their localization dynamics at the stable PGC cytoplasmic bridge.

Our results indicate that ANI-2 is not required for proper P_4_ cytokinesis or for the block of abscission that we observe in this germ line precursor blastomere. This implies that there is a specific, ANI-2-independent mechanism that is active in the P_4_ blastomere to block abscission. Germ cell abscission failure has been characterized in other organisms such as mouse and *Drosophila* and has been shown to promote germ cell syncytial organization (Greenbaum et al., 2006; Huynh et al., 2001). Several contractility regulators that we show localize to the stable PGC cytoplasmic bridge during embryogenesis are also found enriched at stable intercellular bridges connecting germ cells to the rachis in adult *C. elegans* animals (Amini et al., 2014; Rehain-Bell et al., 2017; Zhou et al., 2013), suggesting a functional relationship between the two developmentally-distinct structures. Every adult germ cell in the *C. elegans* gonad appears to be connected to the rachis by an intercellular bridge however, making it unlikely that this adult structure would directly arise from the single stable bridge connecting the two PGCs. Whether the stable PGC cytoplasmic bridge is required later for syncytial organization will require further investigation.

We find that the presence of ANI-2 at the stable cytoplasmic bridge is required around the 200-cell stage to prevent its collapse. This coincides with a reorganization of embryonic tissues during which the two PGCs initiate a rotation along the antero-posterior axis of the embryo (see Fig. 1 and 4). While the exact role of this rotation event is unclear, it is conceivable that the forces enabling rotation of the two PGCs cause mechanical stress, and the presence of ANI-2 at the stable bridge may be required to compensate for this stress and prevent membrane regression. We previously showed that the presence of ANI-2 at stable intercellular bridges in adult germ cells enables them to sustain deformation and that adult *ani-2*(*-*) mutant animals also display a severe membrane collapse defect that begins to manifest itself when oogenic cytoplasmic flows initiate in the gonad, an event that we postulated causes stress on the intercellular bridges (Amini et al., 2014). A balanced regulation between short and long anillin isoforms at cytoplasmic bridges may therefore be a selected molecular mechanism to ensure that they remain stable across a range of conditions that impose mechanical stress and deformation.

## MATERIALS AND METHODS

### *C. elegans* strain maintenance

Animals were cultured as previously described (Brenner, 1974) and grown at 20°C, with the exception of animals bearing temperature sensitive alleles, which were maintained at 15°C. For all experiments, worms were synchronized at the L1 larval stage by hatching embryos in M9 buffer (0.022 mM KH_2_PO_4_, 0.042 mM Na_2_HPO_4_, 0.086 mM NaCl, 1 mM MgSO_4_) after sodium hypochlorite treatment (1.2% NaOCl, 250 mM KOH) and allowed to grow on OP50 bacteria to the desired developmental stage. Strains used in this study are listed in Table S1.

### RNAi-mediated depletions

To prepare RNAi plates, stationary phase cultures of HT115 bacteria transformed with individual clones from the Ahringer library (all clones were verified by sequencing, see Table S2 for details; (Kamath et al., 2003)) were diluted 1:60 and grown at 37°C for ∼2.5-3hrs, to an OD_600_ of 0.4-0.6 (log phase). One mM IPTG was added to the cultures prior to seeding onto Nematode Growth Medium plates containing 1 mM IPTG and 100 μg/ml carbenicillin, to a final concentration of 2 mM. RNAi plates were left to dry for 24 hrs. Adult animals were added to plates and depletion time-courses were performed by feeding each RNAi clone in Table S2 and monitoring phenotypes every 2 hours for 48 hrs to find conditions in which a protein of interest was depleted enough to see an effect on embryonic viability (∼50% hatching failure), cytokinesis (as observed by binucleation of some somatic cells) and still allow the division of P_4_ into Z_2_ and Z_3_ (Fig. S3A). In each experiment, an RNAi clone containing empty vector was used as control.

RNAi clones targeting the genes *rho-1, ptc-1* and *ptc-2* could not be recovered from the Ahringer library and were thus constructed by amplifying the cDNA of each gene by PCR (using the specific primer pairs designed in (Sonnichsen et al., 2005), cloning the PCR product in vector L4440 and transforming ligated products into E. coli strain HT115. All clones were verified by sequencing. Primer pairs were as follows: *rho-1* forward: 5′-AAAAAGATATCATCGTCTGCGTCCACTCTCT-3′, *rho-1* reverse: 5′ - AAGGCTCGAGCTCGGCTGAAATTTCCAAAA - 3′; *ptc-1* forward: 5′ - AAGAAAGATCTGATCGAATCTGCTGGTTGTG - 3′, *ptc-1* reverse: 5′ - AAAAAGCTTCATTTTTCGAGAGAGCTGGC - 3′; *ptc-2* forward: 5′ - AAGAAAGATCTGAGATTCAAGCAGAGCCTGG -3′, *ptc-2* reverse: 5′ - AAAAAGCTTGAGCACAGAATGATCGCAGA - 3′.

### Live imaging

Embryos were obtained by cutting open gravid hermaphrodites in M9 buffer using two 25-gauge needles and mounted on a poly-L-lysine-coated coverslip. The coverslip was flipped on a 3% agarose pad on a glass slide and sealed with valap (vaseline:lanolin:paraffin, 1:1:1). To image P_4_ cytokinesis and midbody release, 2-4 cell-stage embryos were manually sorted before mounting and incubated at 20°C for 5 hr until the embryos had reached the desired developmental stage (*i.e*. P_4_ before division). Images were acquired with a 2x2- or 3x3-binned Axiocam 506 camera (Zeiss) camera mounted on a Zeiss Observer Z1 inverted microscope equipped with a Yokogawa CSU-X1 spinning disk confocal head and illuminated with 488 nm and 561 nm laser lines controlled by Zen software (Zeiss). Embryos were visualized with either a 40×/1.3 NA or 63×/1.4 NA Plan Apochromat oil objective and z-stacks (0.5 μm or 0.75 μm) sectioning the entire embryo were acquired every 35 sec for 2 hr (Fig. 2 or Fig. 3A-B) or every 5 min for 5 hr (Fig. 3 C-D).

To visualize cytoplasmic bridge integrity, embryos were mounted as above and images were acquired with a Nikon A1R laser-scanning confocal microscope and illuminated in single-track mode with 488 nm and 561 nm laser lines controlled by NIS Elements 4.2 software (Nikon). Embryos were visualized with an Apo 40×/1.25 NA water-immersion objective and z-stacks (0.75 μm) sectioning the entire embryo were acquired every 3 min for 3-5 hr. Images were processed and analyzed from the original files using ImageJ software.

### Indirect immunofluorescence

Following RNAi depletions, embryos were collected by hypochlorite treatment, washed in M9 buffer and deposited in 15 μL of M9 on a 14 × 14-mm patterned Cel-Line slide (Thermo Fisher Scientific) coated with 0.1% poly-l-lysine. A coverslip was placed on top of the embryos and the slide immediately put for 30 min on a pre-chilled metal block on dry ice. The coverslip was flipped to crack the egg-shell and the slide immediately immersed in -20°C methanol for 20 min. The slide was allowed to dry and the sample was rehydrated in phosphate buffer saline (PBS, 137 mM NaCl, 2.7 mM KCl, 10 mM Na_2_HPO_4_, 1.8 mM KH_2_PO_4_) for 5 min and then thrice with PBS containing 0.05% Tween 20 (PBST) for 5 min each, and 30 min in blocking buffer (PBST, 10% donkey serum). To diminish background due to non-specific antibody interactions, primary and secondary antibodies were added sequentially. Samples were first incubated overnight at 4°C in 25 μL of PBST with primary antibodies [rabbit anti-ANI-2 (1:1000; (Maddox et al., 2005)) and mouse anti-P granules (clone OIC1D4, 1:300; (Strome, 1986)). After three washes of 5 min in PBST, secondary antibodies [Alexa Fluor 568–coupled donkey anti–rabbit and Alexa Fluor 647–coupled donkey anti–mouse (1:500 each; Invitrogen)] were applied for 90 min at room temperature followed by three washes in PBST. Samples were then incubated for 90 min at room temperature in 25 μL of PBST with goat anti-GFP antibodies (1:1000; Rockland), followed by three PBST washes and incubation with Alexa Fluor 488-coupled donkey-anti-goat for another 90 minutes at room temperature. Samples were first washed for 20 minutes with PBST containing DAPI (1 μg/ml) and two additional PBST washed. Samples were mounted in Prolong Gold (Invitrogen) and allowed to dry overnight at room temperature before being imaged. Images were acquired with a Nikon A1R laser-scanning confocal microscope and illuminated in single-track mode with 405 nm, 488 nm, 561 nm and 640 nm laser lines controlled by NIS Elements 4.2 software (Nikon). Embryos were visualized with a 63×/1.4 NA Plan Apochromat oil objective and z-stacks (0.5 μm) sectioning the entire embryo were acquired. Images were processed and analyzed from the original files using ImageJ software.

Temperature-sensitive mutant embryos were processed as above except for a few differences in how they were initially collected and processed. They were first collected and washed in solutions at 15°C before being divided in two populations. Each population was suspended in M9 buffer at either 15°C (permissive, control) or 30°C (restrictive) and placed for 5 min in 15°C and 25°C incubators, respectively. Embryos were collected by centrifugation (for 1.5 min) at the permissive or restrictive temperature and immediately processed as above for freeze cracking, fixation, staining and imaging. Hence, the total treatment time at restrictive temperature was for a minimum of 5 min (initial incubation) and maximum of 10 min (subsequent processing time). This experimental setting enabled us to bypass essential gene requirements during P_4_ cytokinesis and monitor phenotypic consequences into PGCs that were born and developed at permissive temperature.

### Quantitative image analysis

To determine the number of nuclei per embryo, confocal z-stacks were 3D reconstructed using Imaris software (version 7.6.5) to obtain voxel signals for an entire embryo. Using the “objects” tool, nuclei were defined and counted as objects in the DAPI (immunofluorescence) or mCherry::H2B (live) channel with a diameter of 2.4 ± 0.6 μm (depending on the embryonic stage).

The timings of cytoplasmic isolation and midbody ring release during cytokinesis were determined from time-lapse images using ImageJ software. For each movie, average projection images of three consecutive Z planes were generated and fluorescence intensities of the membrane marker and tagged contractility proteins was measured along a 3 pixel-wide line drawn parallel (cytoplasmic isolation) or perpendicular (midbody release) to the cleavage furrow (see Fig. S1). Fluorescence background was measured in a different region of the cell and subtracted from the furrow measurements. Cytoplasmic isolation was ascribed as the first time point when two ingressing peaks of fluorescence intensity coalesced into a single peak (apparent membrane closure). Midbody ring release was ascribed as the first time point when the peak of fluorescence intensity for a tagged contractility regulator became distant by more than 0.4 μm from the membrane marker at the cell boundary. Cytoplasmic bridge collapse was ascribed as the first time point when the continuous membrane fluorescence signal between the two PGCs separated into two distinct peaks of intensity (apparent membrane collapse).

The focus volume and sphericity of proteins that stably localize between Z_2_ and Z_3_ was done on images from fixed or live samples using Imaris software. For each set of experiment, the control (RNAi) condition was used to set up the baseline values to do the 3D reconstruction of the cytoplasmic bridge foci as an isosurface (see Fig. 4B), using the “surface tool”. First, the intensity of the 3D rendering of the fluorescence signal (either ANI-2 or GFP) was adjusted to get an appropriate signal-to-noise ratio to visualize the focus between Z_2_ and Z_3_. Then using the “surface” tool, a cubic region was drawn around the Z_2_ and Z_3_ cells (identified using the germ granules signal) and an isosurface was automatically generated (using a surface “grain size” set to 0.1 μm) to obtain the 3D reconstruction. The isosurface was then adjusted manually (by changing the threshold values, which were kept thereafter for all foci quantified in that set of experiment) so both the rendering and the reconstruction would fit into one shape. Then, a filter was applied to exclude of the 3D reconstruction all 3D renderings having 10 voxels or less. Once reconstructed, the values given for the reconstruction’s volume (μm^3^) were used as markers of ANI-2 or GFP-tagged protein accumulation between Z_2_ and Z_3_. The values of area (μm^2^) were also used to measure sphericity (Ψ), for which the relationship between focus volume (V) and its surface area (A) was determined using the equation Ψ = π^1/3^(6V)^2/3^/A (Wadell, 1935).

### Statistical analysis

GraphPad software was used for graphing and all statistical analyses. When assumptions of normality and equal variance were met for the data analyzed (Figs. 1, 2, 3), a parametric test of statistical significance between samples was performed by applying a Student’s t-test (one comparison) or a one-way analysis-of-variance (ANOVA) followed by a *post hoc* Dunnett’s test (two or more comparisons). In cases when these assumptions failed to be met (Figs. 5, 6, 7), non-parametric tests were used (Kruskal-Wallis with Dunn’s as *post hoc* for multiple comparisons). In all cases a two-tailed *p*-value smaller than 0.05 was considered significant. All results are expressed as average ± standard deviation (SD) unless otherwise indicated. Sample size (*n*) is given in each figure panel or legend. All results shown are representative of at least three independent biological replicates for each condition, except for Fig. 5 (N≥2). In all figures, *p*-values are represented as follow: *p<0.05, **p<0.01, ***p<0.001, and n.s. (not significant) is p>0.05.

## ACKNOWLEDGEMENTS

We thank Jim Priess, Julie Canman, Karen Oegema, Michael Glotzer, Abby Gerhold and Amy Maddox for strains and reagents, and Nicolas Chartier and Gilles Hickson for comments on the manuscript. We are also grateful to Christian Charbonneau of IRIC’s Bio-imaging Facility for technical assistance and all members of the Labbé laboratory for helpful discussions. Some strains were provided by the CGC, which is funded by NIH Office of Research Infrastructure Programs (P40 OD010440). E.G. is a research fellow of the Fonds de la Recherche du Québec - Santé (FRQ-S). R. A. is a research fellow of the Natural Sciences and Engineering Research Council of Canada (NSERC). This study was supported by NSERC grant #434942 to J.-C. L. IRIC is supported in part by the Canada Foundation for Innovation and the FRQ-S.

## AUTHORS CONTRIBUTION

E. G., R. A. and J.-C. L. designed the experiments. E. G. and R. A. performed all experiments and analyzed data. E.G. and J.-C.L. wrote the manuscript with inputs from R.A.

## REFERENCES

Agromayor, M., and J. Martin-Serrano. 2013. Knowing when to cut and run: mechanisms that control cytokinetic abscission. Trends Cell Biol. 23: 433–441.

Amini, R., E. Goupil, S. Labella, M. Zetka, A.S. Maddox, J.C. Labbé, and N.T. Chartier. 2014. *C. elegans* Anillin proteins regulate intercellular bridge stability and germline syncytial organization. J Cell Biol. 206: 129–143.

Brenner, S. 1974. The genetics of *Caenorhabditis elegans*. Genetics. 77: 71–94.

Carlton, J.G., A. Caballe, M. Agromayor, M. Kloc, and J. Martin-Serrano. 2012. ESCRT-III governs the Aurora B-mediated abscission checkpoint through CHMP4C. Science. 336: 220–225.

Chartier, N.T., D.P. Salazar Ospina, L. Benkemoun, M. Mayer, S.W. Grill, A.S. Maddox, and J.C. Labbé. 2011. PAR-4/LKB1 mobilizes nonmuscle myosin through anillin to regulate *C. elegans* embryonic polarization and cytokinesis. Curr Biol. 21: 259–269.

Chisholm, A.D., and J. Hardin. 2005. Epidermal morphogenesis. WormBook:1–22.

D’Avino, P.P., M.G. Giansanti, and M. Petronczki. 2015. Cytokinesis in animal cells. Cold Spring Harb Perspect Biol. 7: a015834.

D’Avino, P.P., T. Takeda, L. Capalbo, W. Zhang, K.S. Lilley, E.D. Laue, and D.M. Glover. 2008. Interaction between Anillin and RacGAP50C connects the actomyosin contractile ring with spindle microtubules at the cell division site. J Cell Sci. 121: 1151–1158.

Davies, T., S.N. Jordan, V. Chand, J.A. Sees, K. Laband, A.X. Carvalho, M. Shirasu-Hiza, D.R. Kovar, J. Dumont, and J.C. Canman. 2014. High-resolution temporal analysis reveals a functional timeline for the molecular regulation of cytokinesis. Dev Cell. 30: 209–223.

Fawcett, D.W., S. Ito, and D. Slautterback. 1959. The occurrence of intercellular bridges in groups of cells exhibiting synchronous differentiation. J Biophys Biochem Cytol. 5: 453–460.

Frenette, P., E. Haines, M. Loloyan, M. Kinal, P. Pakarian, and A. Piekny. 2012. An anillin-Ect2 complex stabilizes central spindle microtubules at the cortex during cytokinesis. PLoS One. 7: e34888.

Green, R.A., H.L. Kao, A. Audhya, S. Arur, J.R. Mayers, H.N. Fridolfsson, M. Schulman, S. Schloissnig, S. Niessen, K. Laband, S. Wang, D.A. Starr, A.A. Hyman, T. Schedl, A. Desai, F. Piano, K.C. Gunsalus, and K. Oegema. 2011. A high-resolution *C. elegans* essential gene network based on phenotypic profiling of a complex tissue. Cell. 145: 470–482.

Green, R.A., J.R. Mayers, S. Wang, L. Lewellyn, A. Desai, A. Audhya, and K. Oegema. 2013. The midbody ring scaffolds the abscission machinery in the absence of midbody microtubules. J Cell Biol. 203: 505–520.

Green, R.A., E. Paluch, and K. Oegema. 2012. Cytokinesis in animal cells. Annu Rev Cell Dev Biol. 28: 29–58.

Greenbaum, M.P., L. Ma, and M.M. Matzuk. 2007. Conversion of midbodies into germ cell intercellular bridges. Dev Biol. 305: 389–396.

Greenbaum, M.P., W. Yan, M.H. Wu, Y.N. Lin, J.E. Agno, M. Sharma, R.E. Braun, A. Rajkovic, and M.M. Matzuk. 2006. TEX14 is essential for intercellular bridges and fertility in male mice. Proc Natl Acad Sci U S A. 103: 4982–4987.

Gregory, S.L., S. Ebrahimi, J. Milverton, W.M. Jones, A. Bejsovec, and R. Saint. 2008. Cell division requires a direct link between microtubule-bound RacGAP and Anillin in the contractile ring. Curr Biol. 18: 25–29.

Haglund, K., I.P. Nezis, and H. Stenmark. 2011. Structure and functions of stable intercellular bridges formed by incomplete cytokinesis during development. Commun Integr Biol. 4: 1–9.

Hickson, G.R., and P.H. O’Farrell. 2008. Rho-dependent control of anillin behavior during cytokinesis. J Cell Biol. 180: 285–294.

Hirsh, D., D. Oppenheim, and M. Klass. 1976. Development of the reproductive system of Caenorhabditis elegans. Dev Biol. 49: 200–219.

Hubbard, E.J., and D. Greenstein. 2005. Introduction to the germ line. WormBook:1–4.

Huynh, H., C.Y. Ng, C.K. Ong, K.B. Lim, and T.W. Chan. 2001. Cloning and characterization of a novel pregnancy-induced growth inhibitor in mammary gland. Endocrinology. 142: 3607–3615.

Kachur, T.M., A. Audhya, and D.B. Pilgrim. 2008. UNC-45 is required for NMY-2 contractile function in early embryonic polarity establishment and germline cellularization in *C. elegans*. Dev Biol. 314: 287–299.

Kamath, R.S., A.G. Fraser, Y. Dong, G. Poulin, R. Durbin, M. Gotta, A. Kanapin, N. Le Bot, S. Moreno, M. Sohrmann, D.P. Welchman, P. Zipperlen, and J. Ahringer. 2003. Systematic functional analysis of the Caenorhabditis elegans genome using RNAi. Nature. 421: 231–237.

Kechad, A., S. Jananji, Y. Ruella, and G.R. Hickson. 2012. Anillin acts as a bifunctional linker coordinating midbody ring biogenesis during cytokinesis. Curr Biol. 22: 197–203.

Kuwabara, P.E., M.H. Lee, T. Schedl, and G.S. Jefferis. 2000. A *C. elegans* patched gene, *ptc-1*, functions in germ-line cytokinesis. Genes Dev. 14: 1933–1944.

Liu, J., G.D. Fairn, D.F. Ceccarelli, F. Sicheri, and A. Wilde. 2012. Cleavage furrow organization requires PIP(2)-mediated recruitment of anillin. Curr Biol. 22: 64–69.

Maddox, A.S., B. Habermann, A. Desai, and K. Oegema. 2005. Distinct roles for two *C. elegans* anillins in the gonad and early embryo. Development. 132: 2837–2848.

Mishima, M., S. Kaitna, and M. Glotzer. 2002. Central spindle assembly and cytokinesis require a kinesin-like protein/RhoGAP complex with microtubule bundling activity. Dev Cell. 2: 41–54.

Morita, E., V. Sandrin, H.Y. Chung, S.G. Morham, S.P. Gygi, C.K. Rodesch, and W.I. Sundquist. 2007. Human ESCRT and ALIX proteins interact with proteins of the midbody and function in cytokinesis. EMBO J. 26: 4215–4227.

Mullins, J.M., and J.J. Biesele. 1977. Terminal phase of cytokinesis in D-98s cells. J Cell Biol. 73: 672–684.

Oegema, K., M.S. Savoian, T.J. Mitchison, and C.M. Field. 2000. Functional analysis of a human homologue of the *Drosophila* actin binding protein anillin suggests a role in cytokinesis. J Cell Biol. 150: 539–552.

Pacquelet, A., P. Uhart, J.P. Tassan, and G. Michaux. 2015. PAR-4 and anillin regulate myosin to coordinate spindle and furrow position during asymmetric division. J Cell Biol. 210: 1085–1099.

Piekny, A.J., and M. Glotzer. 2008. Anillin is a scaffold protein that links RhoA, actin, and myosin during cytokinesis. Curr Biol. 18: 30–36.

Rehain-Bell, K., A. Love, M.E. Werner, I. MacLeod, J.R. Yates, 3rd and A.S. Maddox. 2017. A Sterile 20 Family Kinase and Its Co-factor CCM-3 Regulate Contractile Ring Proteins on Germline Intercellular Bridges. Curr Biol. 27: 860–867.

Sonnichsen, B., L.B. Koski, A. Walsh, P. Marschall, B. Neumann, M. Brehm, A.M. Alleaume, J. Artelt, P. Bettencourt, E. Cassin, M. Hewitson, C. Holz, M. Khan, S. Lazik, C. Martin, B. Nitzsche, M. Ruer, J. Stamford, M. Winzi, R. Heinkel, M. Roder, J. Finell, H. Hantsch, S.J. Jones, M. Jones, F. Piano, K.C. Gunsalus, K. Oegema, P. Gonczy, A. Coulson, A.A. Hyman, and C.J. Echeverri. 2005. Full-genome RNAi profiling of early embryogenesis in *Caenorhabditis elegans*. Nature. 434: 462–469.

Straight, A.F., C.M. Field, and T.J. Mitchison. 2005. Anillin binds nonmuscle myosin II and regulates the contractile ring. Mol Biol Cell. 16: 193–201.

Strome, S. 1986. Asymmetric movements of cytoplasmic components in *Caenorhabditis elegans* zygotes. J Embryol Exp Morphol. 97 Suppl:15–29.

Sulston, J.E., E. Schierenberg, J.G. White, and J.N. Thomson. 1983. The embryonic cell lineage of the nematode *Caenorhabditis elegans*. Dev Biol. 100: 64–119.

Sun, L., R. Guan, I.J. Lee, Y. Liu, M. Chen, J. Wang, J.Q. Wu, and Z. Chen. 2015. Mechanistic insights into the anchorage of the contractile ring by anillin and Mid1. Dev Cell. 33: 413–426.

Wadell, H. 1935. Volume, Shape, and Roundness of Quartz Particles. The Journal of Geology. 43: 250–280.

Wang, J.T., and G. Seydoux. 2013. Germ cell specification. Advances in experimental medicine and biology. 757: 17–39.

Wheatley, S.P., and Y. Wang. 1996. Midzone microtubule bundles are continuously required for cytokinesis in cultured epithelial cells. J Cell Biol. 135: 981–989.

Zhou, K., M.M. Rolls, and W. Hanna-Rose. 2013. A postmitotic function and distinct localization mechanism for centralspindlin at a stable intercellular bridge. Dev Biol. 376: 13–22.

